# Dynamics of Human Serum Transferrin in Varying Physicochemical Conditions Explored by Using Molecular Dynamics Simulations

**DOI:** 10.1101/2022.01.28.478130

**Authors:** Sowmya Indrakumar, Alina Kulakova, Pernille Harris, Günther H. J. Peters

**Author notes:** Biomolecular Sciences, University of Copenhagen, 2100 Copenhagen, Denmark (Sowmya Indrakumar). Department of Chemistry, University of Copenhagen, 2100 Copenhagen, Denmark (Alina Kulakova, Pernille Harris).

## Abstract

Conformational stability of human serum transferrin (Tf) at varying pH, salt, and excipient concentrations were investigated using molecular dynamics (MD) simulations and the results are compared with previously published small angle X-ray scattering (SAXS) experiments. SAXS study showed that at pH 5, Tf is predominantly present in partially open (PO) form, and the factions of PO differ based on the physicochemical condition and drifts towards closed form (HO) as the pH increases. Tf is a bilobal glycoprotein that is composed of homologous halves termed as N-lobe and C-lobe. The current study shows that the protonation of Y188 and K206 at pH 5 is the primary conformational drive into PO, which shifts towards the closed (HO) conformer as the pH increases. Furthermore, at pH 6.5, PO is unfavorable due to negative charge-charge repulsion at the N/C-lobe interface linker region causing increased hinge distance when compared to HO, which has favorable attractive electrostatics. Subsequently, the effect of salt concentration at 70 and 140 mM NaCl was studied. At 70 mM NaCl and pH 5, chloride ions bind strongly in the N-lobe iron-binding site, whereas these interactions are weak at pH 6.5. With increasing salt concentration at pH 5, regions surrounding the N-lobe iron-binding site are saturated and as a consequence sodium and chloride ions accumulate into the bulk. Additionally, protein-excipient interactions were investigated. At pH 5, excipients interact in similar loop regions, E89-T93, D416-D420, located in the C-lobe and N-lobe of the HO conformer, respectively. It is anticipated that interactions of additives in these two loop regions cause conformational changes that lead to iron coordinating residues in the N-lobe to drift away from iron and thus drive HO to PO conversion. Furthermore, at pH 6.5 and 140 mM histidine or phosphate, these interactions are negligible leading to the stabilization of HO.

## INTRODUCTION

Transferins are a family of proteins that belong to the group of non-heme ferric ion (Fe^3+^) binding glycoproteins widely present in bodily fluids and cells of vertebrates.[1–3] Transferrin (Tf) which is synthesized in the liver[4] and secreted into the plasma, is the main ferric ion transporter. Tf delivers iron to target sites like the liver, the spleen, and bone marrow where it is incorporated into newly formed erythrocytes via the transferrin receptor (TfR) mediated endocytosis.[5] During endocytosis at acidic pH, the entire Tf-TfR complex is internalized into the endosome to release ferric ion and subsequently, the complex is returned to the cell surface. Tf is then released from the complex back into the cytosol to acquire more iron.[6] Unbound ferric ions participate in redox reactions (e.g. Fenton reaction) to produce free radicals that are known to attack proteins, nucleic acids, etc. Therefore, the binding of ferric ions to Tf is of prime importance to creating an anti-bacterial, and low-iron environment.[7] Tf is an 80 kDa bilobal glycoprotein (half-life 8-10 days) with two homologous halves termed as N-lobe and C-lobe, respectively, and each lobe binds to a ferric ion.[1] The presence of anions such as bicarbonate (BCT)[8] facilitates the binding of iron by excluding water from the coordination sites otherwise the binding is negligible.[2]

Tf’s potential in a pharmaceutical application has been established. Tf supplements are recommended to treat disease conditions such as anemia and toxicity conditions of iron such as oxidative stress.[9] Apart from Fe^3+^, Tf has been shown to bind to other metal ions such as Ga^3+^, In^3+^, Ti^4+,^ and Ru^3+^ for therapeutic or diagnostic applications.[10–12]

Moreover, the efficient cellular uptake pathway has been explored for targeted delivery of small-molecule drugs, or other macromolecules by Tf.[11,13] Transferrin-conjugated drugs have shown significant improvement in cytotoxicity and selectivity of the drugs.[14] Moreover, the efficacy of the Tf-conjugated drug as an anticancer therapy has been demonstrated in the *in-vitro* treatment of breast cancer cells and prostate cancer cells in mouse models.[15] Additionally, Tf-conjugated drugs can cross the blood-brain barrier by receptor-mediated endocytosis targeting rat tumor cells, but not human cells posing further formulation challenges. [15–18] One of the major limitations in the development of protein therapeutics is the proper stabilization of biologics as these structures are dynamic and may have sites prone to chemical (oxidation and hydrolysis) and physical (aggregation and unfolding) degradation.[19] To overcome these challenges, excipients are used in an effort to stabilize biologics without losing their biological activity and to prevent protein-protein interactions that might cause protein instability[20–22] and consequently lead to insoluble aggregates. The general formulation approach is to screen for different physicochemical conditions such as pH, excipients, and buffer systems to improve protein stability by for instance varying pH and/or osmolality of the solution.[23–29] To optimize protein formulation, a molecular understanding of excipient-protein interactions is pivotal and can guide scientists in designing formulations.

Several studies have been conducted on Tf in the line of pH and salt effects on protein properties.[30–32] For instance, a study showed that change in the protonation state of Y188 at low pH initiates iron release from the N-lobe.[30,31,33] Another study focusing on the salt effect on protein properties showed that salt accumulation starts first in the N-lobe, and iron release from the N-lobe is coordinated by N-lobe and C-lobe interactions.[32,34] To the best of our knowledge, there exists no computational study which discusses excipient and salt effects as a function of pH on Tf stability. To investigate the molecular origin for the preference of the different conformers at different physicochemical conditions as deduced from small angle X-ray scattering (SAXS), molecular dynamics (MD) simulations were utilized. The different physicochemical conditions were chosen based on an earlier SAXS study, which demonstrated that at low pH partially open (PO) conformer is preferred and the equilibrium shifts towards the closed conformer (HO) as the pH increases. [24] At low pH, the fraction of PO differs depending on salt and excipient concentrations. In the current study, we have performed MD simulations to obtain a detailed molecular understanding of how these physicochemical conditions affect protein surface properties and conformational stability, which can be utilized in designing preventive measures of aggregation in optimizing protein formulations.

## METHODOLOGY

### Structure

Several X-ray crystal structures of Tf are available, and they represent three different conformations, i.e. iron-bound diferric holo form (PDB ID: 3V83)[35], iron-free apo form (PDB ID: 2HAV)[6], and intermediate form termed as “partially open (PDB ID: 3QYT)[36] with N-lobe open as seen in apo form and C-lobe closed as seen in the holo form with Fe^3+^ bound in the two lobes. The partially open and holo-form of Tf will be referred to as PO and HO, respectively, in the following. Figure 1 shows the structural architecture of Tf and residues coordinating with Fe^3+^ and BCT in the two lobes. Each lobe has two subdomains. The N-lobe consists of N’(1-95 and 247-331) and N’’(96-246). The C-lobe comprises C’(339-425 and 573-679) and C’’(426-572). A linker (331-339) connects both lobes. The Tf structure is stabilized by 19 cystines, and the C-lobe has an extra cystine making it more rigid compared to the N-lobe.

**Figure 1.**
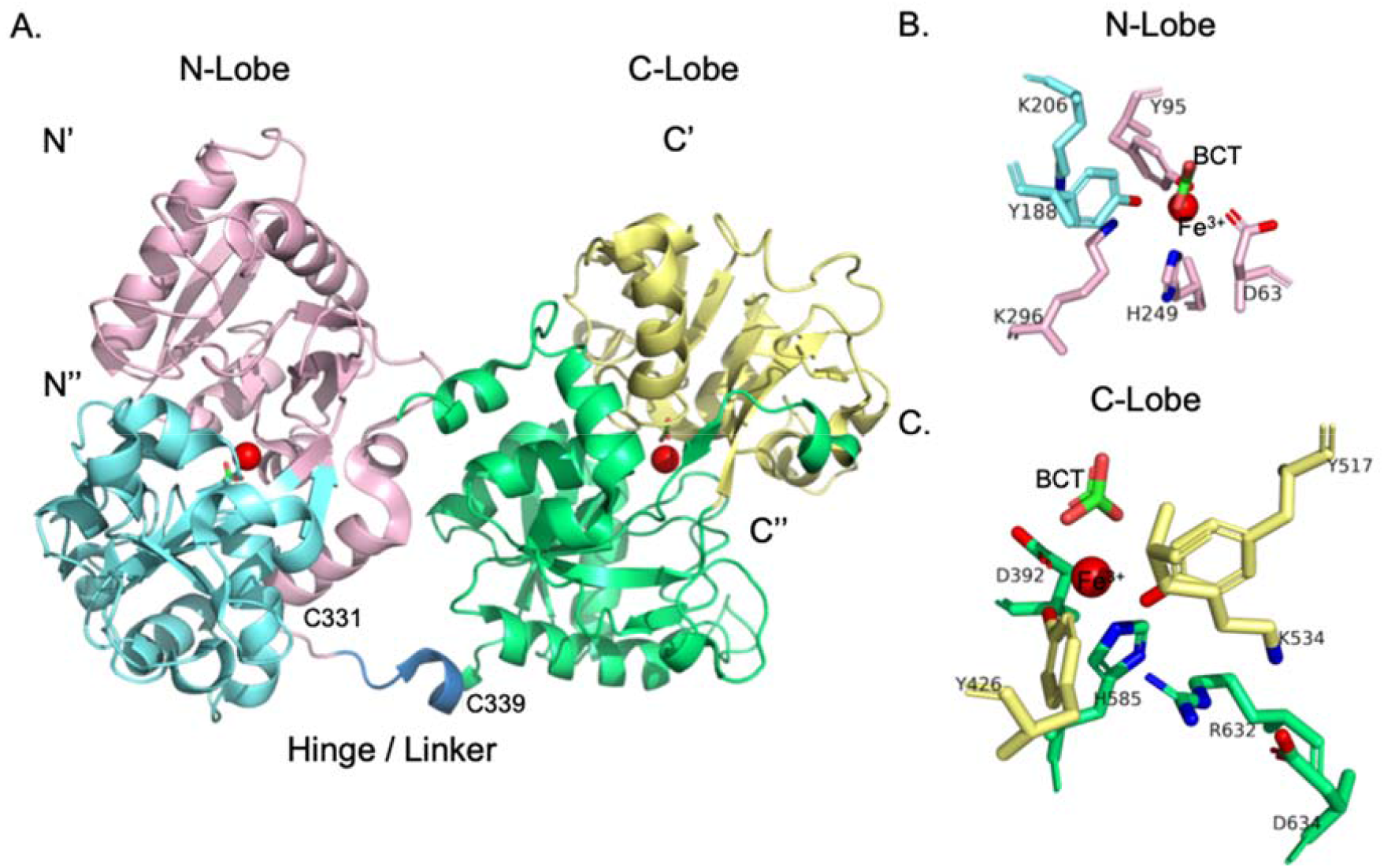
Structure of Tf. Color coding is as follows: pink (N’), cyan (N’’), green (C’), yellow (C’’), and blue (linker residues 331-339)[7]. Fe^3+^ ions are shown as red spheres, and BCT is displayed in a stick representation, where oxygen and carbon atoms are colored in red and green, respectively. B. Coordination of the residues and BCT molecule with Fe^3+^ in the N-lobe is displayed. Residues D63, Y95, Y188, and H249 coordinate with Fe^3+^, and a dilysine bridge also termed as ‘dilysine trigger’[37] (K206, K296) constitutes the second shell residues. C. Coordination of the residues and BCT molecule with Fe^3+^ in the C-lobe is depicted. Residues D392, Y426, Y517, and H585 coordinate with Fe^3+^, and the triad of residues, K534, R632, and D634, forms the second shell residues.

### Molecular dynamics simulations

The different conformations of transferrin were subjected to MD simulations at different physicochemical conditions to investigate the conformational changes induced by pH change and the addition of salt and excipients. To ease the discussion, salt ions and excipients will also be referred to as additives.

#### 1. Effect of pH

The protonation state of the titratable amino acids at pH values of 5, 6.5, and 8 was calculated using the H++ server (http://biophysics.cs.vt.edu/H++).[38] The server settings were adjusted to an external dielectric constant of 80 for water, and the internal dielectric constant was set to 10.[38] The topology and coordinate files generated as output in Amber format were used to generate the PDB structure file using ambpdb tool implemented in Amber 16.[39] The PDB files were used as the starting structure for all-atom classical constant pH MD simulations[40] in explicit solvent. Simulations were performed with the Amber 16 program[41] employing the amber force field, ff99SB[42], for proteins. Water molecules were represented using the TIP3P[43] water model. Parameters for Fe^3+^ were defined using the 12-6-4 LJ-type nonbonded[44,45] model in the amber force field. The bicarbonate molecule was prepared using antechamber[46] module in Amber 16 and the AM1-BCC[47] charge method. All bonds to hydrogen atoms were constrained using the SHAKE algorithm.[48] Each system was neutralized using chloride ions (pH 5, 6.5, and 8 needed 8, 5, and 1 chloride ion(s), respectively). Each system was solvated in a truncated octahedron water box with a 15 Å padding in all three Cartesian directions. The particle mesh Ewald method[49] was employed to determine the non-bonded electrostatic energies with a real-space cutoff of 8 Å. Each system was minimized for 5000 steps. The first 1500 steps were performed using the steepest descent method followed by the conjugate gradient method for the remaining steps. Subsequently, each system was heated linearly from 10 K to 300 K within 1ns using the Langevin thermostat[50] with a collision frequency of 5 ps^−1^. Berendsen barostat[51] was used to control pressure dynamics. The systems were then equilibrated for 2 ns at constant temperature (300 K) and pressure (1 bar). Following this, a short equilibration of 2 ns at constant pH was performed to update the protonation of titratable residues. Only, the residues near Fe^3+^ were titrated during the simulation to take into account fluctuation in the protonation states. Finally, constant pH simulations were performed for 100 ns, and coordinates were saved every 10 ps. Each system at a particular pH was simulated in duplicates for 100 ns to estimate the statistical uncertainty of the results. Different starting structures for the simulations were obtained by applying different steepest descent minimization cycles (1500 and 2000 steps).

#### 2. Effect of Salt

The effect of 70 and 140 mM NaCl was studied at pH 5 and 6.5. The ionic strength (IS) was adjusted to 70 and 140 mM NaCl by the addition of 126 (Na^+^ + Cl^-^), and 248 (Na^+^ + Cl^-^) ions, respectively, to the solvated system containing ∼48000 water molecules. Each system at a particular physiochemical condition of pH and IS was simulated in duplicates for 100 ns as described above.

#### 3. Effect of excipients

Arginine, acetate, phosphate, and histidine were considered since those were used previously in the SAXS study on Tf stability.[24] Structures were obtained from PubChem[52] and Zinc Database[53]. These molecules were prepared at the desired pH using the ligprep tool in the Schrödinger suite 2016-3 (Schrödinger, LLC, New York, NY, USA)[54]. Parameter files for the excipients were prepared at desired pH using the antechamber[46] module in Amber 16 in combination with the AM1-BCC[47] charge method. The concentration of excipients was adjusted to 140 mM to match the conditions of the previous SAXS study on Tf stability.[24] Each system was simulated for 100 ns in duplicates at the same conditions as SAXS.

Different parameters that include preferential interaction coefficient (PIC) and contacts formed between excipients and protein were determined to understand at the molecular level the conformational preferences for PO or HO in the presence of excipients and varying salt concentrations and to map potential sites on the protein surface where salt ions and excipients might bind.

##### Preferential interaction coefficient and interaction hotspot region

Preferential interaction coefficient (*Γ*_23_) was used as a measure of preferential interactions between the additives and protein. *Γ*_23_ defines the overall accumulation of additives on the protein surface, which is defined as follows[55]

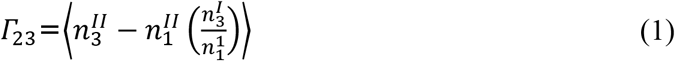

where superscript I represent a bulk region, which is outside of the protein vicinity. Superscript II represents a local region around the protein surface. Angled brackets ⟨ ⟩ denote an ensemble average, and *n* denotes the number of specific additives. Subscripts 1, 2, and 3 stand for water, protein, and additives, respectively. Positive *Γ*_23_ implies favorable protein-excipient interactions, and *vice-versa* when *Γ*_23_ is negative. A cutoff of 8Å was used to differentiate local from the bulk region.[56] To estimate *Γ*_23_, the trajectories from the production runs were divided into 5 ns time intervals each containing 50 frames. An average *Γ*_23_ was calculated for each 5 ns time interval, and uncertainties in *Γ*_23_ are reported as the standard error of the means.[56]

## RESULTS AND DISCUSSION

### 1. Effect of pH

Tf releases ferric ions at endosomal pH 5.6 by opening the iron-binding site.[57] Previously performed SAXS experiments identified different proportions of PO and HO conformers depending on the pH.[24] The equilibrium between HO and PO conformations shifts towards the PO conformer with decreasing pH. At pH 5, ∼55% PO and ∼23% HO were present, whereas, this equilibrium shifts towards the HO conformer (90%) as the pH was increased to 6.5 and above. Several factors such as N-lobe and C-lobe interactions and change in protonation states of iron coordinating residues influence the conformational changes that lead to ferric ion release from the lobe.[30,58] Studies carried out previously have shown that the electrostatic field around Fe^3+^ is important for the opening of the lobes.[59] Therefore, the electrostatic potential surface was calculated using the H++ server[38] considering the most representative structure extracted from clustering of the MD trajectory. Using agglomerative hierarchical clustering approach, the conformations generated over time were grouped into distinct clusters depending on the conformational similarity.[60]

At pH 5, the electrostatic potential surfaces of PO and HO were similar (Figure 2A, B). However, the titratable residues (D63, Y95, Y188, K206, H249, K296, D392, Y426, Y517, K534, H585) coordinating with Fe^3+^ were susceptible to changes in their protonation states (Figure 1). Change in the protonation state of these residues is known to initiate the conformational change since the corresponding residues lose the ability to coordinate to iron.[30,31,58] The simulations reveal that at pH 5, the protonation state of Y95 and Y188 fluctuate between being protonated and deprotonated (on the hydroxyl group) during the 100 ns simulation. During the course of the simulations, Y188 drifted away from the iron, indicating that Y188 is involved in ferric ion release from the N-lobe (Figure S1). This is in line with previous studies which indicated that the opening of the N-lobe is triggered by the protonation of Y188.[61] Further, K206, which is part of the “dilysine trigger”, is protonated most of the time during the simulation at pH 5, whereas, it remains deprotonated at pH 6.5 and 8. Previously, it has been shown that protonation of one of the lysine residues constituting the “dilysine trigger” initiates conformational change for N-lobe opening[62]. Additionally, root mean square fluctuations (RMSFs) of the lobes are slightly higher at pH 5 as compared to pH 8 (Figure 3). The N-lobe has higher fluctuation compared to the C-lobe (Figure 3) owing to the presence of one extra cystine in the C-lobe. The N-lobe fluctuation is rather global; however, we see that local fluctuations are higher around the first shell iron coordinating residues, which may initialize iron release from N-lobe.

**Figure 2.**
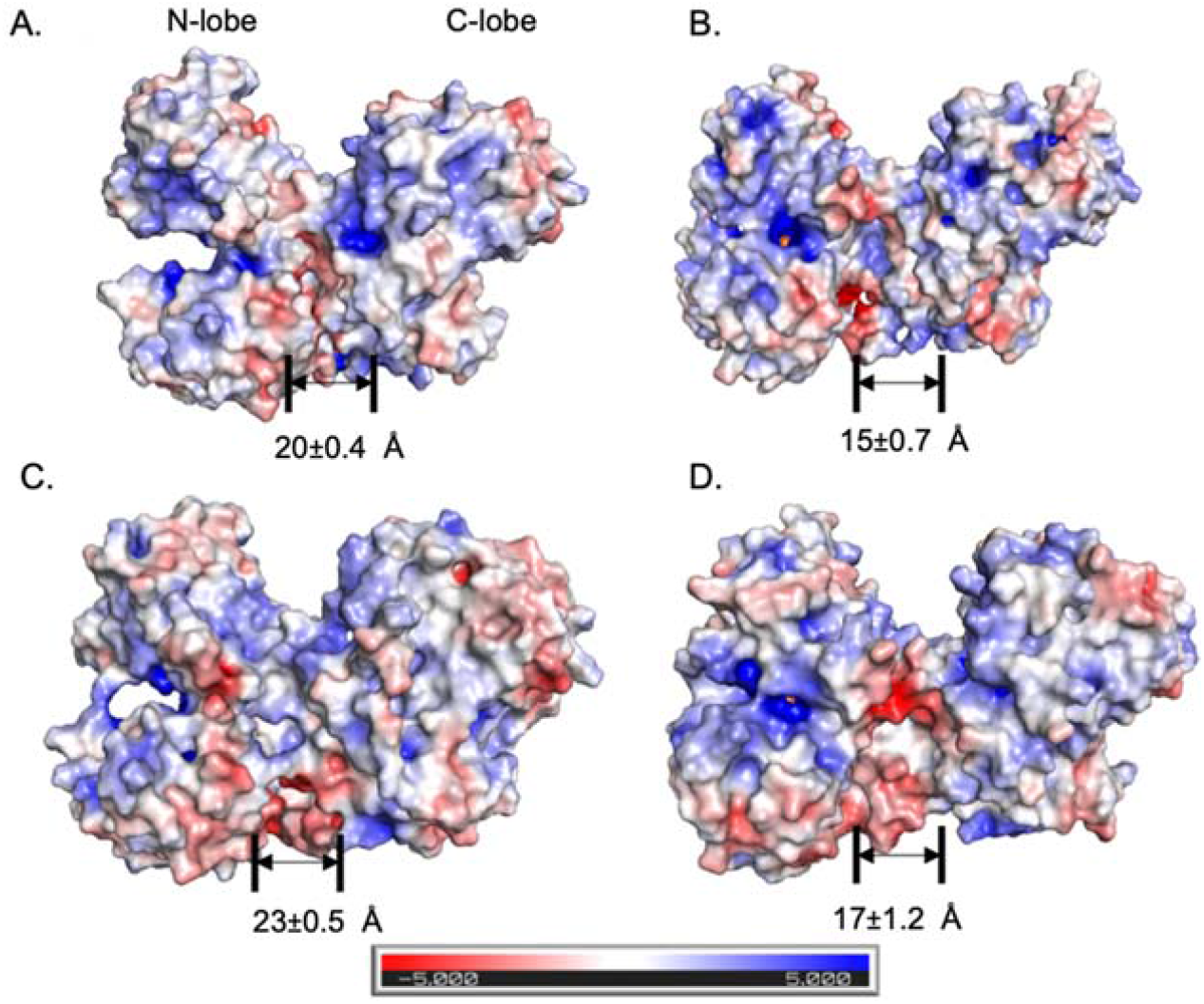
Electrostatic potential surfaces calculated for A. PO, pH 5; B. HO, pH 5; C. PO, pH 6.5; D. HO, pH 6.5. H++ server was used to generate electrostatic potential surfaces. Red, white, and blue surface colors indicate negative, close to neutral and positive potentials, respectively, as indicated by the color bar. The structures are presentative structures extracted from clustering of the MD trajectory using the agglomerative hierarchical clustering approach.[60] Linker (331-339) distances are shown below each image.

**Figure 3.**
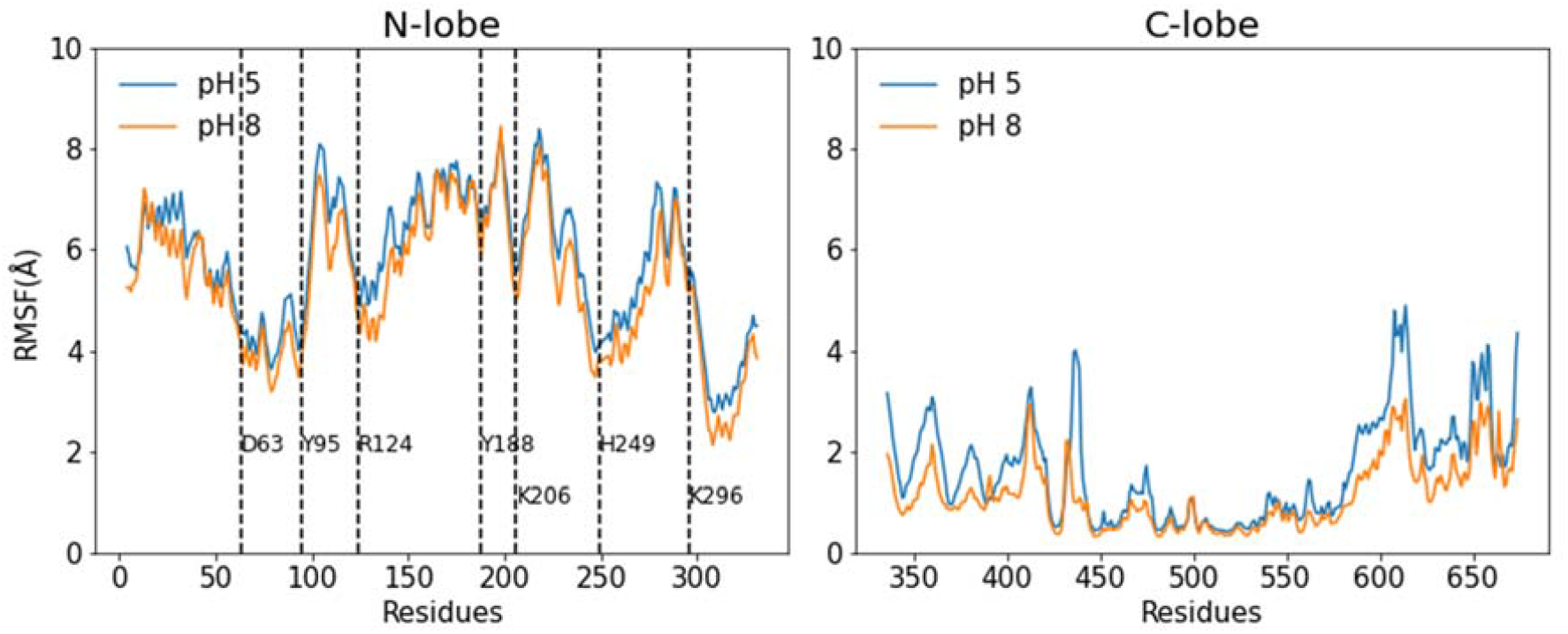
RMSF plots of N-lobe and C-lobe of HO at pH 5.0 (blue line) and 8.0 (orange line) from the combined analysis of the replicas. N-lobe iron coordinating residues are marked with black dashed lines.

Ideally, at physiological pH, Tf exists in the closed (HO) form. During the simulation of the HO at pH 6.5 and 8, neither Y95 nor Y188 was protonated, and Lys206 was deprotonated, keeping the dilysine-bridge intact and no drastic change in conformation was encountered. Root mean square deviation (RMSD) of Tf fluctuated within 4Å (Figure S2). These findings were in agreement with the SAXS results wherein HO conformation existed in high proportion at pH 6.5 and 8.[24] This further supports the view that HO conformation is preferred since Y188 and K206 are deprotonated in the HO conformer. Additionally, at pH 6.5, electrostatics surface plots of the representative PO conformer extracted from the simulations indicated that PO is unfavorable due to negative charge-charge repulsion at the interface linker region (Figure 2C), which leads to an increase in the hinge distance (end-to-end linker residues 331-339) to 23+/-0.5 Å. On the other hand, the interface appeared less positive in the HO conformation with an average hinge distance of 17+/-1.2 Å at pH 6.5 (Figure 2D). Overall, pH affects the preference for different conformations. As discussed above, PO conformation is preferred at pH 5, and with an increase in pH, the electrostatic surface potential changes causing that the Tf structural preference to shift towards HO conformation.

### 2. Effect of salt

Comparative electrostatic surface studies of the PO conformer at pH 5 for varying NaCl concentrations (0, 70 mM, and 140mM) were carried out. It was observed that the iron-binding cleft in the N-lobe had a balance of positive and negative charges at 0 mM NaCl, as opposed to a positively charged surface in the presence of 70 mM NaCl (Figure 4A, B). Previous studies on the salt effect have shown that binding of chloride ions to Tf has a synergistic effect on the ferric ion release from the lobes.[32,63] Further, R124 has been recognized as a kinetically significant anion binding site that accommodates chloride ions and allosterically communicates with the iron coordinating residues.[64] In our simulation, we see more accumulation of Cl-ions around R124 in PO as suppose to HO at pH 5 (Figure 5B). In this work, negative PIC values for chloride ions imply that the protein surface is saturated with ions and thus prefers to accumulate into the bulk (Figure 5A and S3A). Further, in Figure 5B, we observe that chloride ions make more contacts with iron coordinating residues in both conformers suggesting initiation of iron release. These stabilizing interactions causes iron to drift away from the coordinating residues in the N-lobe, thus causing its release (Figure 7B).

**Figure 4.**
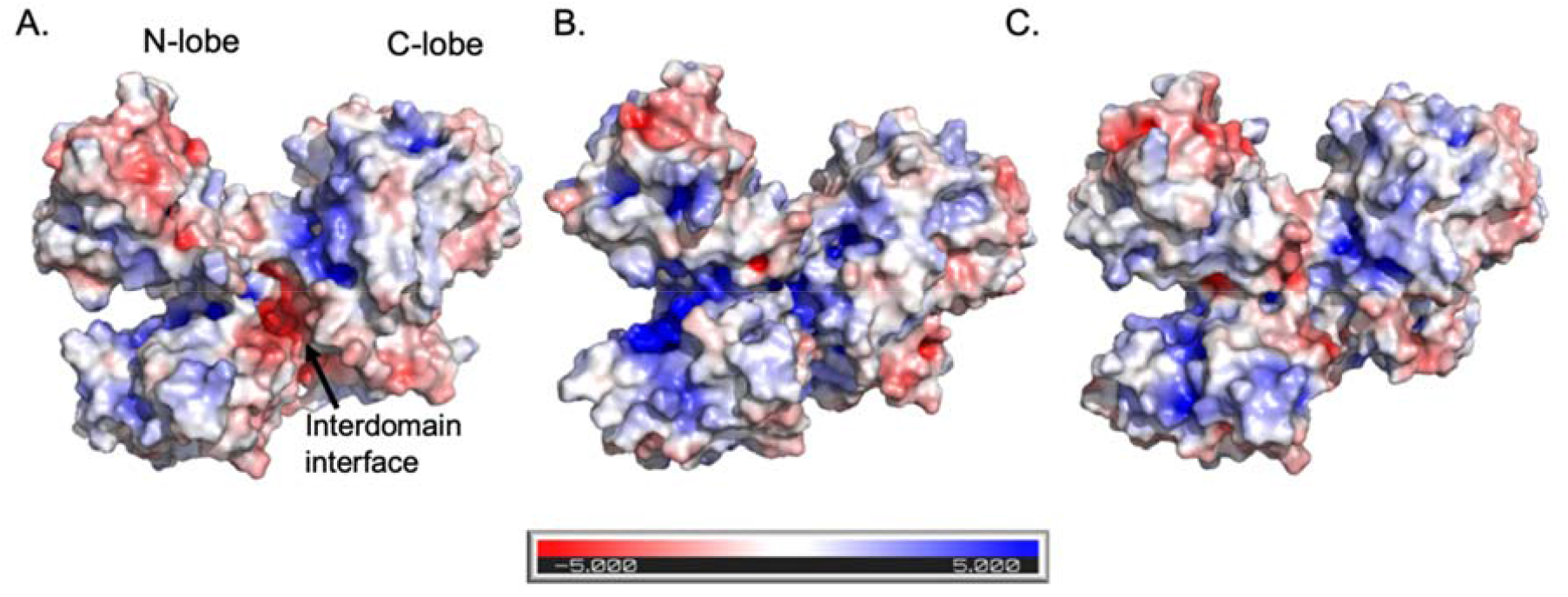
Electrostatics potential surfaces for PO conformation at pH 5 and A. 0 mM NaCl, B. 70 mM NaCl, and C. 140 mM NaCl. The structures are presentative structures extracted from clustering of the MD trajectory using the agglomerative hierarchical clustering approach.[60]

**Figure 5.**
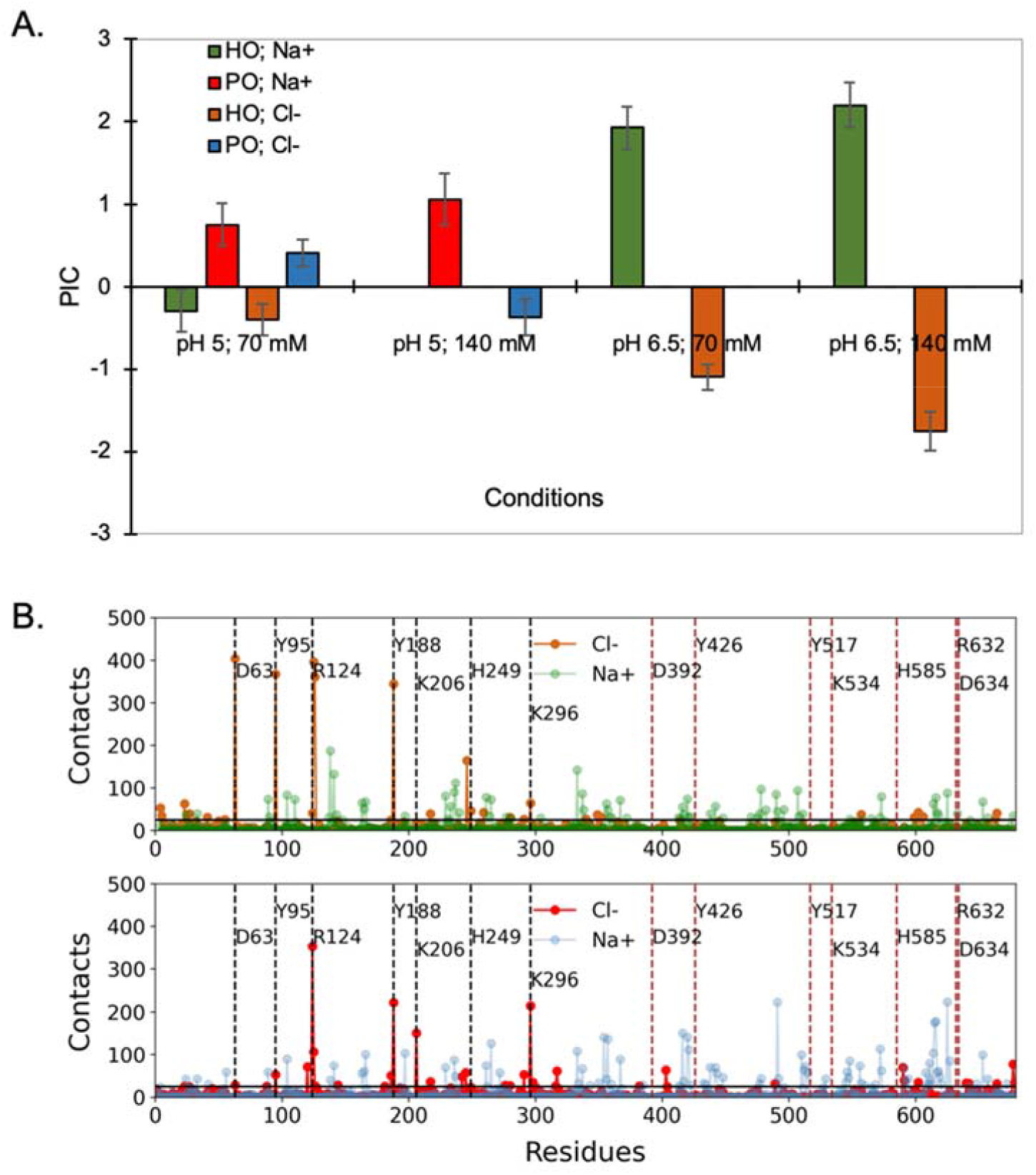
A. PIC values of sodium and chloride ions are shown for the HO and PO conformers at selected conditions corresponding to the conditions used for the SAXS experiments. Positive PIC values imply ongoing salt accumulation on the protein surface or favorable protein-salt interactions while negative values implies saturation of protein surface with salt or unfavorable protein-salt interactions. B. Plot showing the number of contacts formed between ions and HO (top panel) and PO (bottom panel) conformers at pH 5, 70 mM, respectively. The black horizontal line is the standard deviation over all formed contacts across all excipients. N-lobe and C-lobe iron coordinating residues are marked with black and red dashed lines, respectively.

To conclude, as seen in SAXS experiments[24] at pH 5, increasing the salt concentration shifts the equilibrium, PO ⟷ HO, to a higher percentage of PO conformation (from 55% to 64%) reducing the presence of HO conformation (from 23 % to 3%). Further, a strong negatively charged patch around the linker region connecting the N and C-lobes is seen in the absence of salt (Figure 4A). Electrostatics potential around the interdomain linker region shifts from positive to neutral as salt concentration is increased from 70 to 140 mM NaCl (Figure 4B, C), caused by a higher chloride ion accumulation in the interdomain region at 70 mM NaCl as compared to 140 mM NaCl (Figure S3B, C).

For the PO conformer, at pH 5 and 70 mM NaCl, chloride ions strongly interact in the iron-binding cleft of the N-lobe and around the linker region of the protein to balance the positive surface charges in order to stabilize the PO conformer (Figure 5B). At 140 mM NaCl and (pH 5), the PIC values for chloride ions are lower as compared to 70 mM. This implies that saturation of the protein surface with chloride ions has already reached at 70 mM NaCl and a further increase in salt concentration to 140 mM leads to salt accumulation in the bulk (Figure 5A, S3). On the other hand, even though the PIC values for the sodium ions are positive, the absence of sodium ion interaction on the N-lobe iron-binding site imply that sodium may not contribute to the opening of the N-lobe (Figure 5). Unspecific sodium ion binding is observed on the protein surface.

Unlike at pH 6.5, at pH 5, the shift of the equilibrium from HO to PO conformation with the addition of salt can be explained by two effects i.e. change in the protonation states of Y188 and K206; and binding of chloride ions in the iron-binding site of N-lobe (Figure 5B). According to SAXS results, Tf is mainly present in the HO conformation (90%) at pH 6.5, and the addition of NaCl does not promote the PO conformation.[24] At pH 6.5 and 0 mM NaCl, the interface region (around residues D307, E315) is negatively charged (Figure 6A). However, already at 70mM NaCl, there is a balance of positive and negative charges around the interdomain interface region (Figure 6B, C). As reflected by the PIC data (Figure 5A), the trend for chloride and sodium PIC values for HO conformation is similar for the two salt concentrations at pH 6.5. At 70 mM NaCl, there exists chloride interactions with protein around the iron binding site of N-lobe (Y188, R124, H246, K296; Figure S4), though, not as strong as at pH 5 (Figure 5B). Up to the concentration of 140 mM, accumulation of chloride ions is not seen in the iron binding site of N-lobe (Figure S4, top panel). Thus, it shows that the added salt accumulates into the bulk, which is reflected as a high negative PIC value for chloride (Figure 5A, S4). Conversely, sodium ions does not interact with N-lobe iron-binding site but binds unspecifically to the other parts on the protein surface, leading to positive PIC values (Figure 5A, S4). On the whole, HO is stabilized at pH 6.5 by accumulation of salt in the bulk and away from iron binding residues in the N-lobe.

**Figure 6.**
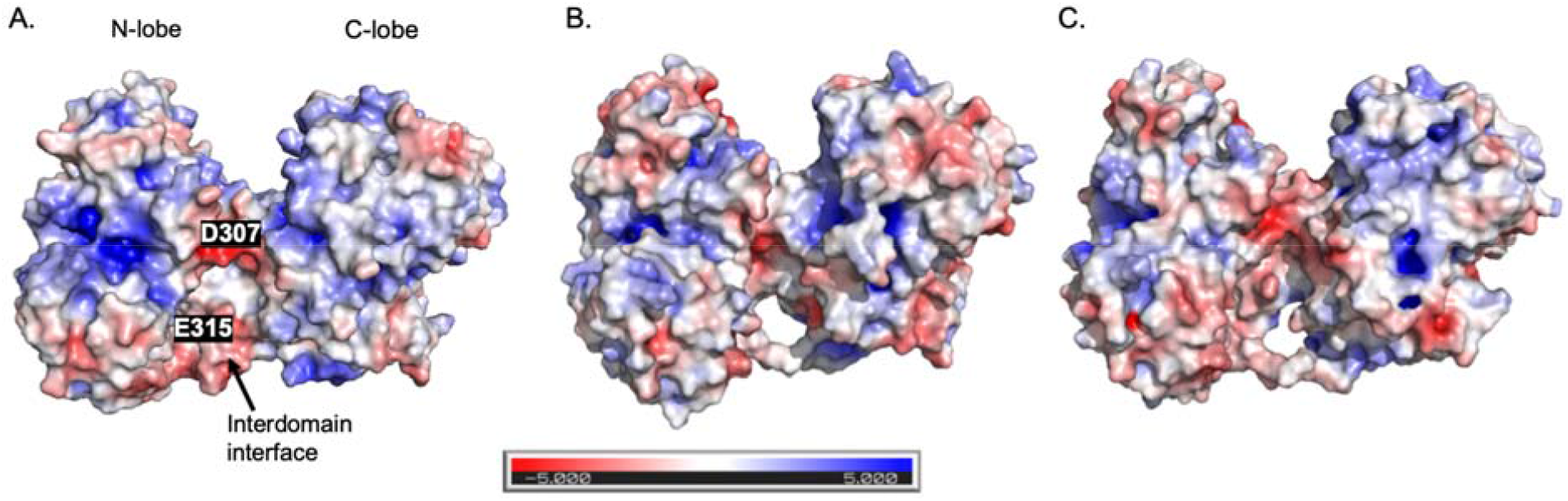
Electrostatic potential surfaces for HO conformation at pH 6.5 and A. 0 mM NaCl, B. 70 mM NaCl and C. 140 mM NaCl. The structures are presentative structures extracted from clustering of the MD trajectory using the agglomerative hierarchical clustering approach.[60]

### 3. Effect of excipients

In the following section, the term excipient is used in a wider range that also includes the buffering agents as excipients[65] since the tested buffer systems not only regulate shifts in pH, but the buffer components also interact with Tf and might affect Tf’s conformational stability. Generally, different conformations are stabilized by the respective excipients by binding to different sites in PO and HO conformers. This is expected due to the different physicochemical properties of the excipients. Besides, at pH 5, arginine and histidine carry a charge of +1 each whereas acetate has a charge of -1. Moreover, arginine has a hydrophobic fatty acid chain and histidine has aromatic properties. Figure 7 depicts the PIC values calculated at selected physicochemical conditions corresponding to the conditions used for the SAXS experiments. The general trend is that similar PICs are observed for arginine and histidine at pH 5 irrespective of the two conformers (HO, PO). Conversely, the negatively charged acetate remains mainly in the bulk phase at pH 5 as indicated by the negative PIC value (Figure 7A). Arginine and histidine make more contacts per residue as suppose to acetate in the HO conformer (Figure 8). At pH 5, distances of the coordinating residues to iron were monitored (based on the center of mass) to see if the addition of excipients can induce conformational changes in Tf (Figure 7B). The increase in the distance in the HO conformer causes iron to drift away from Arg124 to initiate iron release from the N-lobe but no significant change in the Fe^3+^-R124 distance was observed when acetate was added to the solution (Figure 7B,C). However, the distances between iron and Y95/Y188 increased in the presence of acetate and histidine (Figure 7B,C). All in all, the increase in the distance of N-lobe coordinating residues to the iron could cause release of iron.

**Figure 7.**
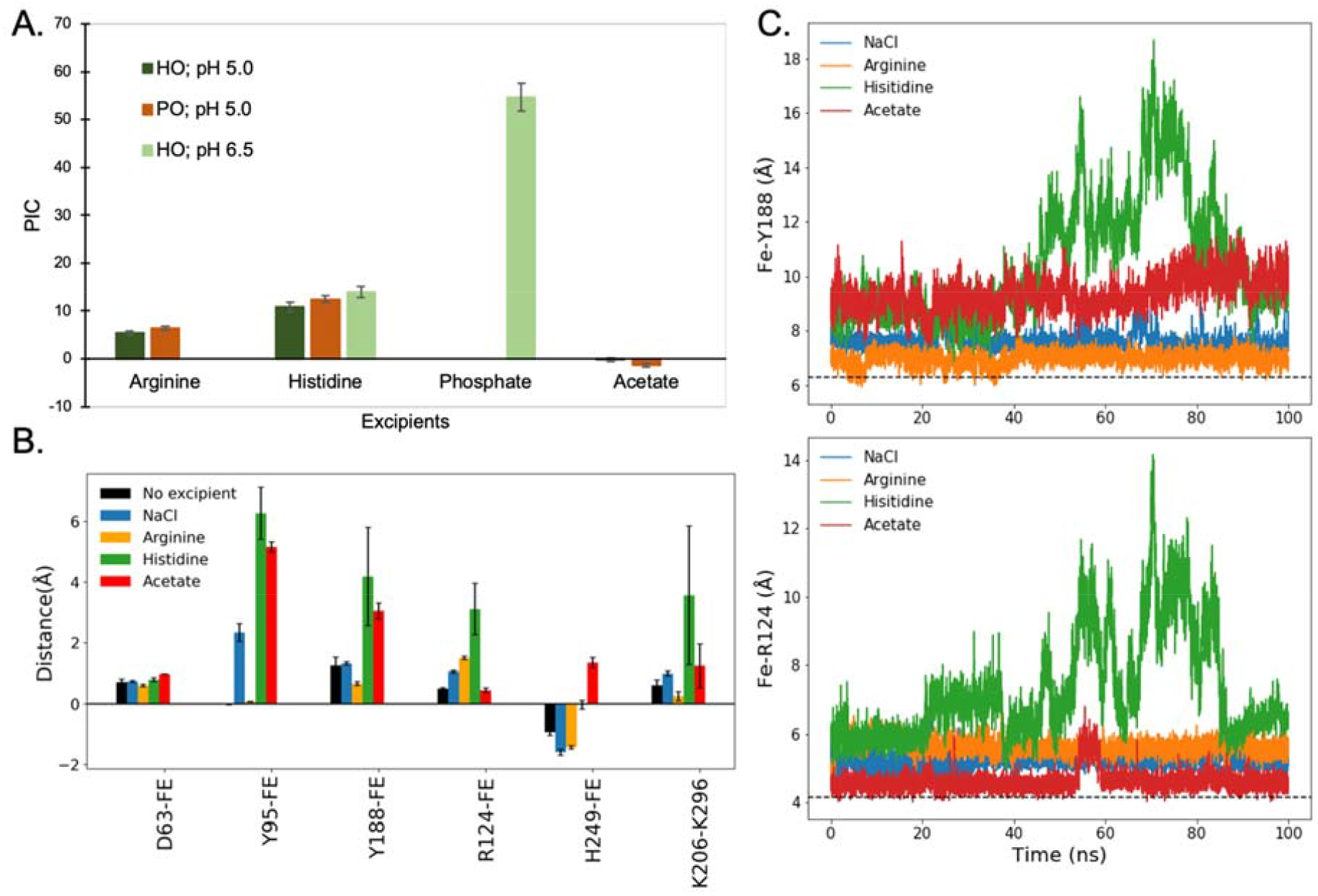
A. *PIC*(*Γ*_23_) values were determined from the simulations of the PO and HO conformers at selected physicochemical conditions corresponding to the SAXS experimental conditions. For phosphate, PIC value is only shown for pH 6.5 in order to compare the PIC values with that of histidine. B. Bar plot showing the difference in the mean distances between iron and the coordinating residues, dislysine bridge for the HO conformer for different excipients at 140mM, pH 5. Mean distances are calculated by subtracting each distance from the respective distance of the minimized starting structure. The error bar is the standard error of the block from the block averaging of the different systems.[66] C. Distance between iron and N-lobe iron coordinating residues (R124, Y188) as a function of time for the HO conformer at pH 5. Black dashed line is the distance in the minimized HO structure.

**Figure 8.**
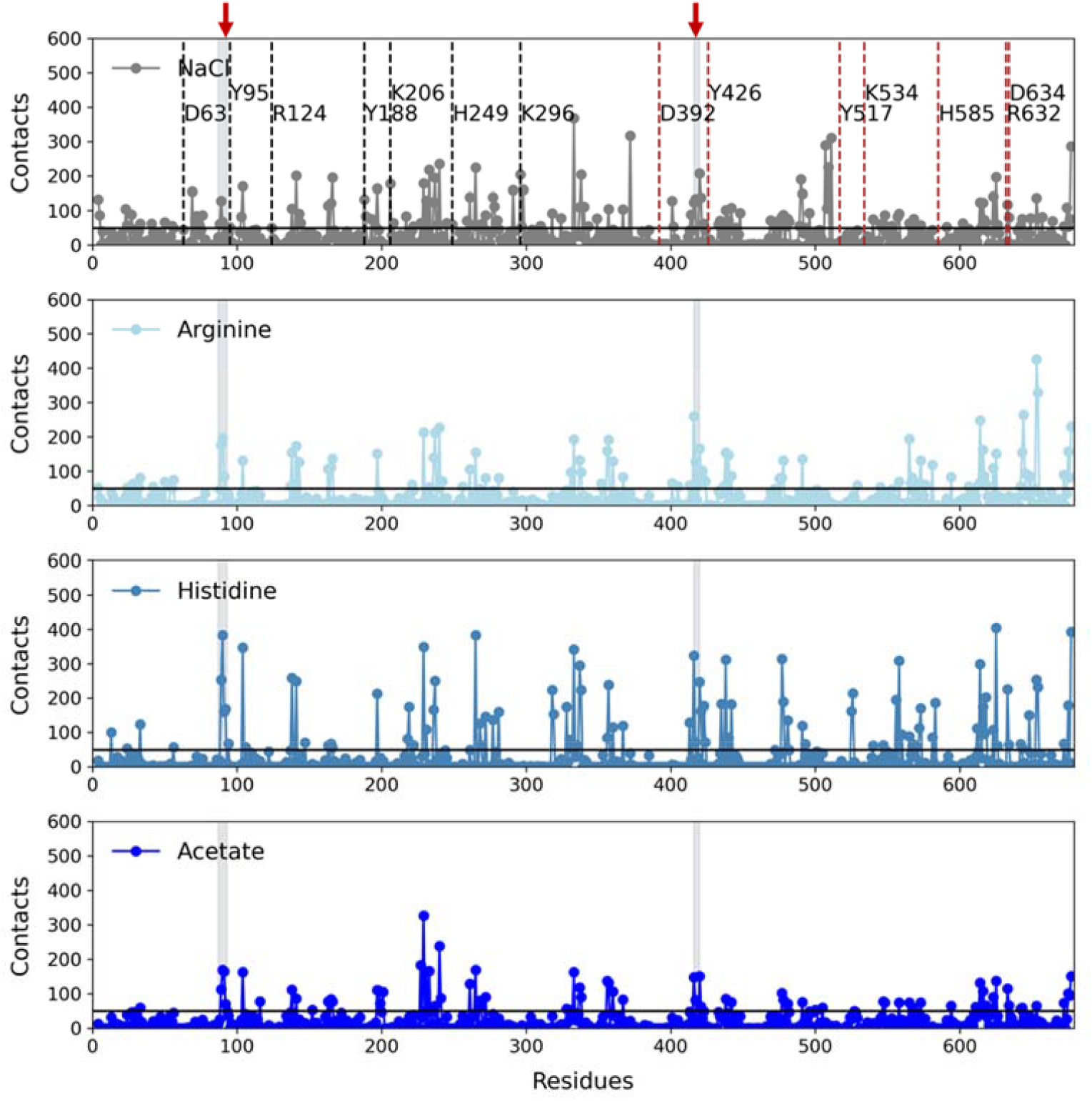
Plot showing number of contacts formed between the excipients and HO conformer at pH 5. Results are shown for excipient concentrations of 140mM. Gray shaded regions with a red arrow are the two loops (89-93, 416-420). The black horizontal line is the standard deviation over contacts considering all four excipients. N-lobe and C-lobe iron coordinating residues are marked with black and red dashed lines, respectively.

Excipients interact with several patches on the protein surface of the HO and PO conformers (Figure 8, S6). Arginine and histidine bind more strongly in the HO and PO conformers as compared to acetate (Figure 8, S6). The differences in the excipient interactions could be the reason for a higher fraction of PO conformation in case of histidine and arginine, but a lower fraction for acetate.[24] However, the common loop regions in the HO conformer where the excipients bind consist of residues D416-D420 (near opening of C-lobe for iron release), and residues E89-T93 (N’-N” connecting loop in N-lobe) (Figure 8). Furthermore, at pH 5 and in the presence of excipients, RMSF is higher depending on the excipient as suppose to in the absence, around the iron binding site in N-lobe including in the two loop regions. This could imply plausible conformational changes to initiate the opening of the N-lobe (Figure S5). These interactions are particularly interesting as they are absent at pH 6.5 for histidine and phosphate (Figure 9). Thus, these specific interactions could possibly initiate conformational changes as reflected in RMSF and change in distances between iron and coordinating residues leading to HO to PO conversion at pH 5 (Figure 7B). In conclusion, the differences in the extent of excipient interactions with Tf could explain the variations in the fraction of PO conformer seen in SAXS experiments.[24]

**Figure 9.**
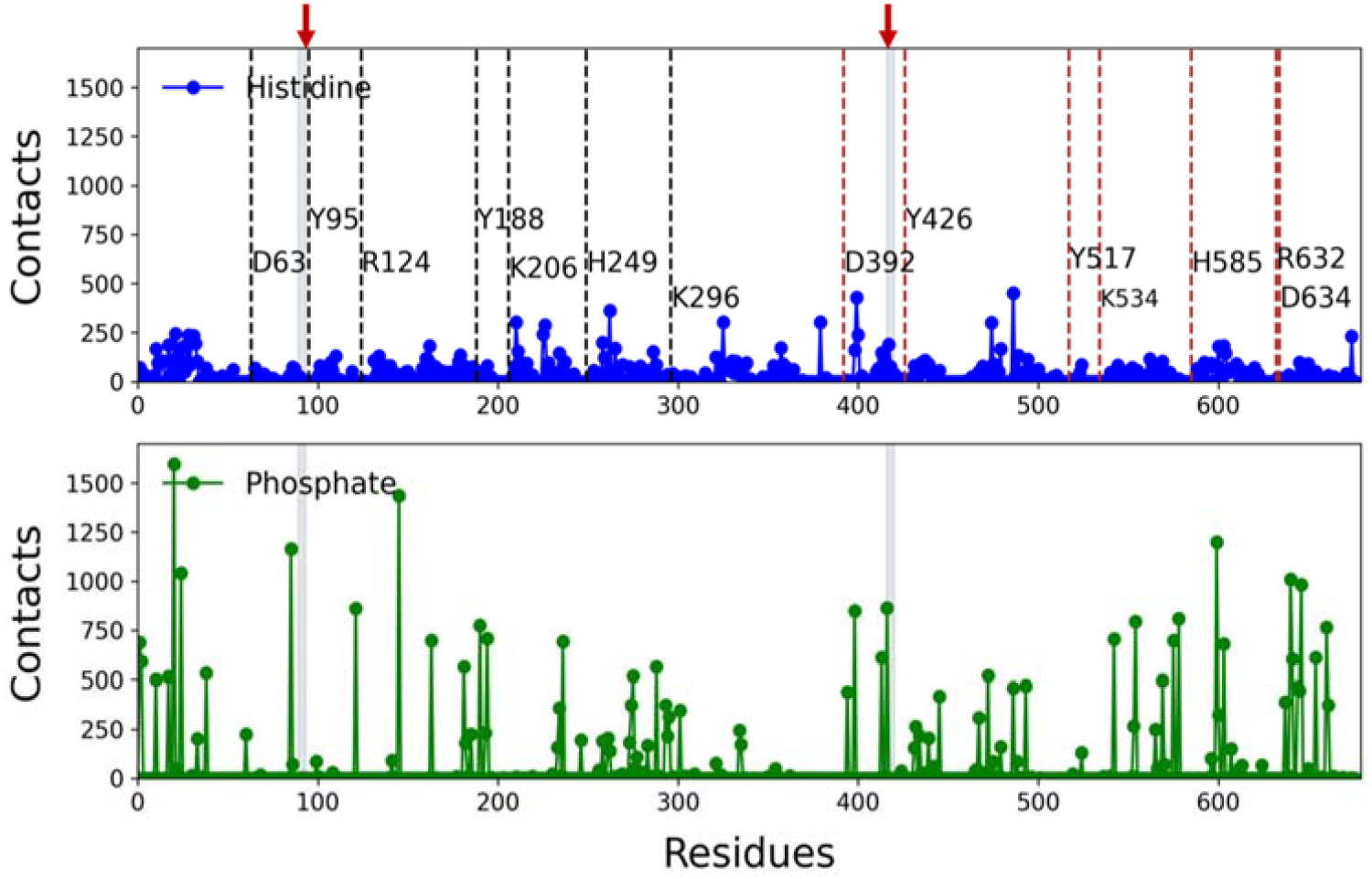
Plot showing number of contacts formed between HO conformer and excipients (Histidine and phosphate on top and bottom panel, respectively) at pH 6.5, 140 mM. Gray shaded regions with a red arrow are the two loops (89-93, 416-420). N-lobe and C-lobe iron coordinating residues are marked with black and red dashed lines, respectively.

At pH 6.5, HO conformation is preferred as Y188 and K206 are deprotonated as discussed in the pH section above. The charge of phosphate and histidine is -2 and 0, respectively, which makes phosphate more likely to interact with the exposed hydrophilic patches strongly as compared to histidine. Phosphate and histidine interactions in the C-lobe loop region (D416-D420) and N-lobe loop region (E89-T93) are negligible at pH 6.5 for the HO conformer (Figure 9). The absence of these interactions could explain why HO conformation is favored at pH 6.5.[24] Returning to the PIC data (Figure 7), phosphate has a substantially higher value compared to other excipients implying a high preference of phosphate for the protein surface. The differences in PIC values can be used to understand the stabilizing ability of excipients.[24] Generally, at pH 6.5, histidine interacts weakly with the protein surface in many regions (Figure 9) causing stabilization of the HO conformer.[24] Interactions for histidine as an excipient in the loop regions (E89-T93, D416-D420) are weaker (Figure 9) than histidine at pH 5 (Figure 8). Conversely, accumulation of phosphate on the protein surface causes destabilization of HO conformer as reflected in RMSF plot and isothermal denaturation experiments (Figure S7).[24]

## CONCLUSIONS

Human serum transferrin (Tf) can exist in three conformers, a holo form[35], an iron-free apo form[6], and an intermediate form called “partially open[36]”. Herein, we have performed MD simulations to investigate the structural properties of the conformers at different physicochemical conditions and the interactions of excipients with the different conformers to map the molecular determinants that determine the preferred conformers observed in SAXS experiments[24]. In this study, the effect of pH 5, 6.5, and 8 were investigated. The main conformational drive into PO conformation at pH 5 is the protonation states of the iron coordinating residues. Previous studies, as well as our studies, have shown that protonation of Y188 brings about change in the protonation of K206 that eventually leads to conformational changes. From SAXS results, it is observed that at pH 5, Tf exists predominantly in the PO conformation, and the equilibrium shifts towards the HO conformer as the pH increases to 6 and above. Our results indicate that the transition between conformers is driven by the change in protonation states of the N-lobe iron coordinating residues (Y188, K206) and a shift in electrostatic surface potential at the interdomain interface. Subsequently, the effect of NaCl at 70 and 140 mM on the lobe dynamics were studied. At pH 5 and 70 mM NaCl, it is encountered that chloride ions bind strongly in the N-lobe iron binding site of HO conformer, whereas these interactions are weak at pH 6.5. The driving force behind the accumulation of the salt in the N-lobe iron binding site is the change in the electrostatic potentials at this region. For 140 mM NaCl, regions surrounding N-lobe iron binding site are saturated and salt accumulates into the bulk away from the protein vicinity. Our results also indicate that depending on the type of excipient conformational changes can be induced. At low pH, arginine and histidine make more contacts with the protein as suppose to acetate due to their different physicochemical properties. All three excipients (arginine, histidine, and acetate) interact in similar loop regions located in the C-lobe (residues D416-D420) and N-lobe (residues E89-T93) of HO conformer. These specific interactions cause conformational changes as reflected in the change in distances between iron and coordinating residues alongside changes in the protonation states of Y188 and K206 thus leading to HO to PO transition. Moreover, the addition of arginine and histidine causes Arg124 in the HO conformation to drift away from the iron. Such a movement is not observed in the presence of acetate suggesting that a larger fraction of PO is present in solution (as seen in SAXS) for arginine or histidine compared to acetate. At pH 6.5, histidine is neutral and interacts only weakly with several patches on the protein surface except at the two loop regions (residues D416-D420 and E89-T93), which is otherwise strong in the two loop regions at pH 5 due to +1 charge thus stabilising PO conformer at low pH. Also, at pH 6.5, phosphate makes more contacts as suppose to histidine due to -2 charge, which might explain the destabilization of of HO conformer and the destabilizing effect as seen in isothermal denaturation experiments.[24] The computational approach presented can be applied to explore conformational dynamics of other biologics.

## AUTHOR CONTRIBUTIONS

S.I. performed the simulations and drafted the manuscript, G.H.J.P. contributed to the design of the computational study and discussion of the results. A.K. and P.H. contributed to the design of SAXS experiments and discussion of the SAXS results. All authors contributed to the writing of the manuscript.

## ACKNOWLEDGMENTS

This study was funded by a project part of the EU Horizon 2020 Research and Innovation program under the Marie Skłodowska-Curie grant agreement No 675074 – “Protein-excipient Interactions and Protein-Protein Interactions in formulation” (PIPPI); http://www.pippi.kemi.dtu.dk. We used VMD 1.9.3 [67], pymol 1.8.4.2[68] and jupyter notebook plugins[69] for performing analyses or making the graphical images. Simulations were performed on the CPU/GPU cluster at DTU Chemistry and the High Performance Computing cluster at DTU. DanScatt is acknowledged for funding the SAXS trip. EMBL P12 DESY and EMBL B29 ESRF are acknowledged for providing beam time for performing the SAXS experiments, and Albumedix Ltd is acknowledged for kindly providing recombinant transferrin.

## ABBREVIATIONS

MD: molecular dynamics
Tf: human serum transferrin
PO: partially open form of Tf
HO: closed form of Tf
AP: open form of Tf
PIC: preferential interaction coefficients
SAXS: small angle X-ray scattering
RMSF: root mean square fluctuation
RMSD: root mean square deviation
TfR: transferrin receptor
BCT: bicarbonate
PDB: protein data bank

## Supplementary material

**Figure S1.**
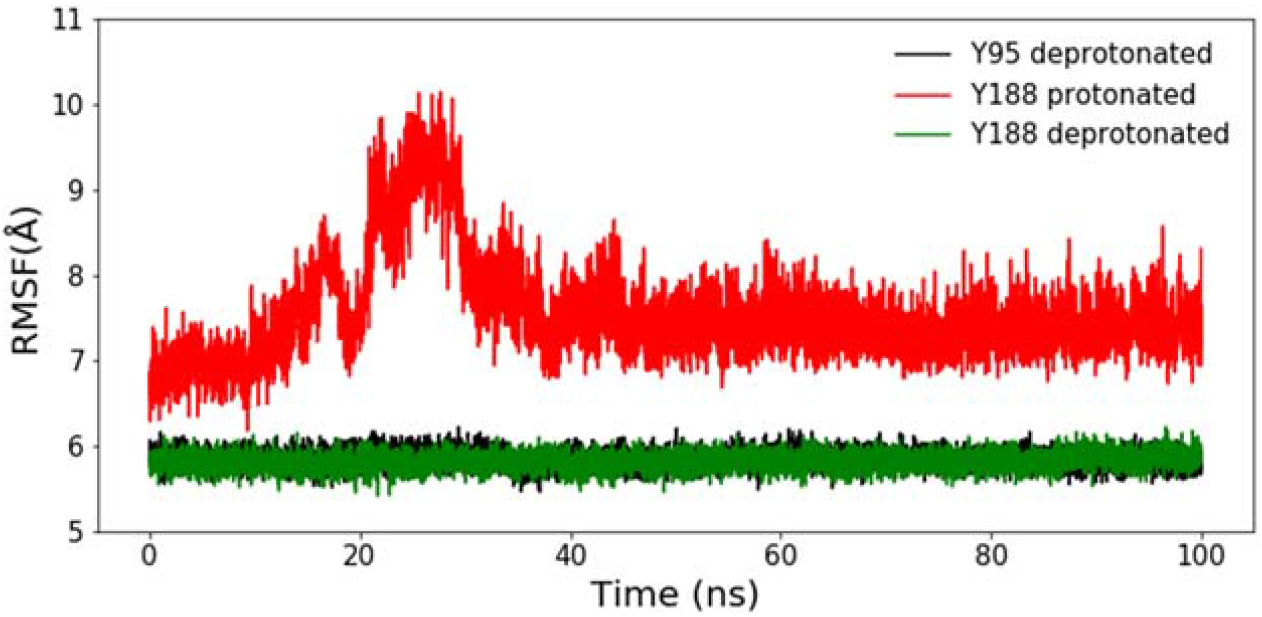
Distance between iron and coordinating Tyr residues at pH 5.0 for HO conformation. Y188 and Y95 were titrated during the simulations, and only Y188 moves away from iron due its protonation (red curve), whereas Y95 stays close to iron (black curve). Additionally, as a test case in another independent simulation, Y188 was kept deprotonated (fixed; green curve) throughout the simulation; it shows that Y188 remains close to the iron (green curve). This demonstrates the role of protonation of Y188 in iron release at pH 5.

**Figure S2.**
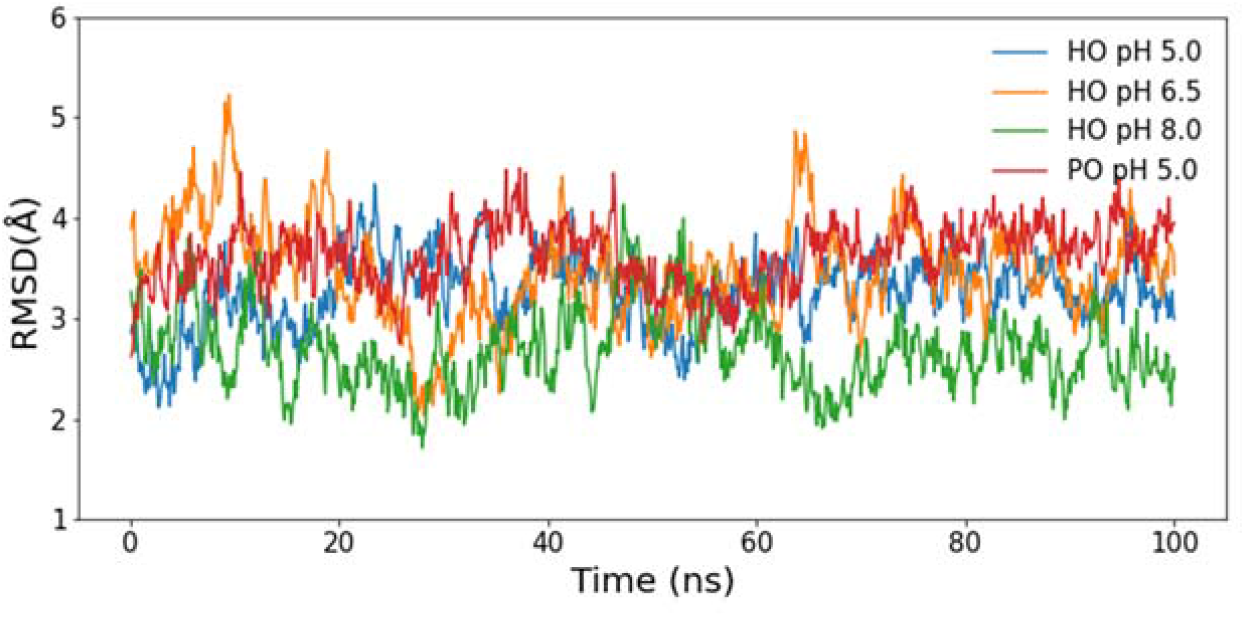
RMSD plot for the different simulated systems.

**Figure S3.**
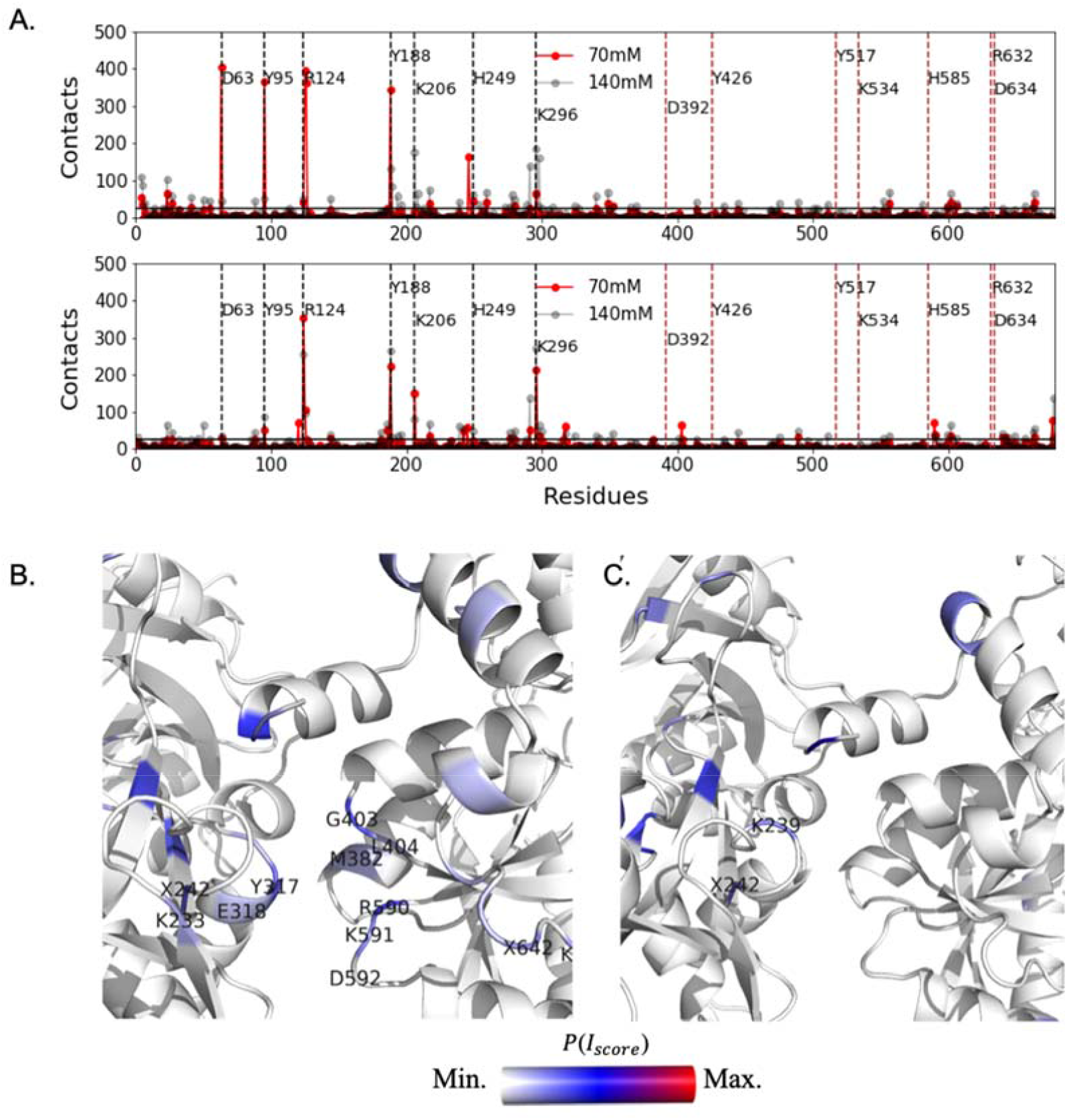
A. Plot showing number of contacts formed between chloride ions and HO (top panel) and PO (bottom panel) conformers at pH 5, 70 mM and 140mM, respectively. Black horizontal line is the standard deviation over contacts determined for HO and PO at the two salt conditions. N-lobe and C-lobe iron coordinating residues are marked with black and red dashed lines, respectively. B. Structure showing subdomain interface of PO conformation colored based on the number of contacts formed with chloride ions at pH 5, 70 mM NaCl. C. Structure of PO conformation colored based on the number of contacts formed with chloride ions at pH 5, 140 mM NaCl.

**Figure S4.**
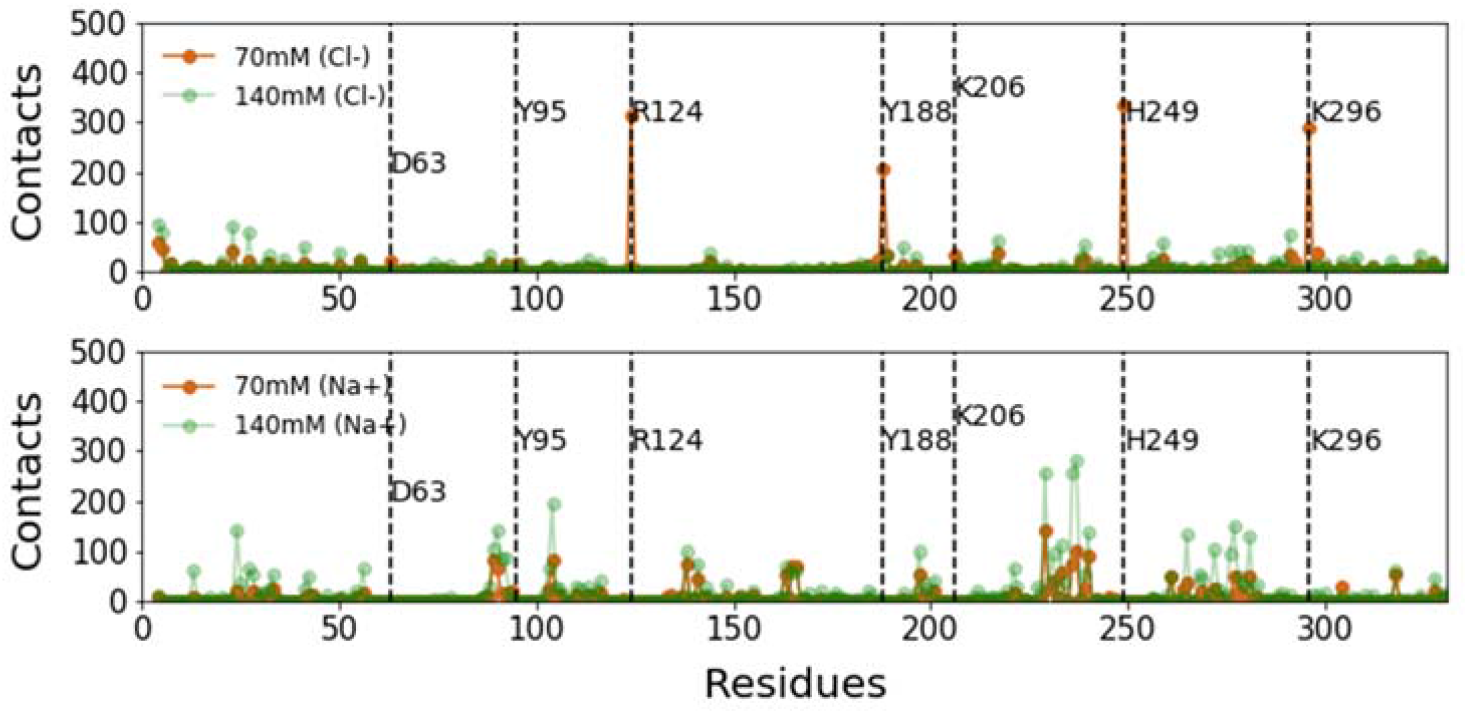
Plot showing number of contacts formed at pH 6.5 between HO conformer and 70 mM and 140mM chloride ions (top panel) and 70 mM and 140 mM sodium ions (bottom). N-lobe iron coordinating residues are marked with black dashed lines.

**Figure S5.**
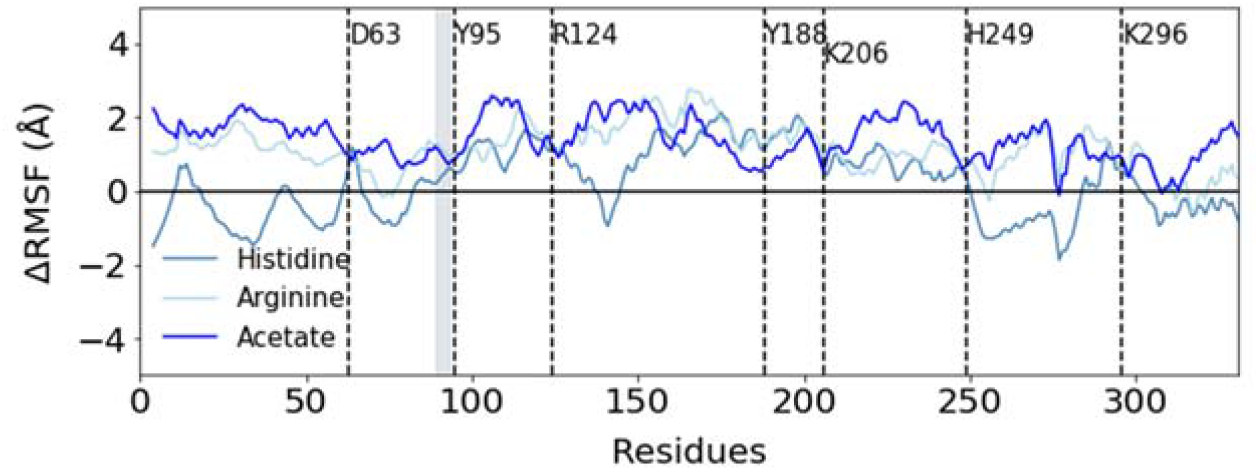
ΔRMSF plot for the different excipient system at pH 5 for HO conformer. RMSF for the different system is subtracted from the system simulated without excipient. Gray shaded regions are the two loops (89-93, 416-420).

**Figure S6.**
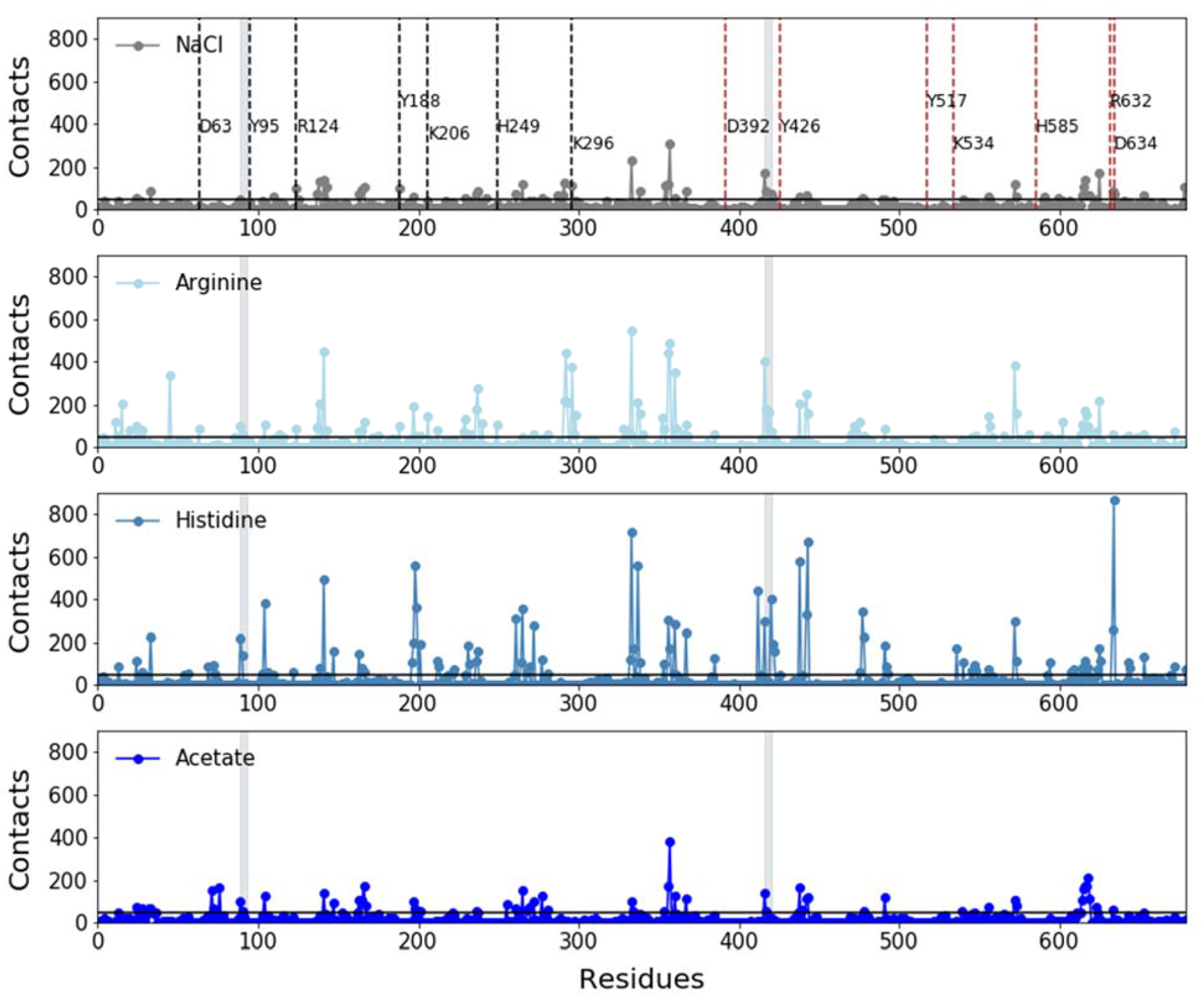
Plot showing number of contacts formed between the excipients and PO conformer at pH 5. Results are shown for excipient concentration of 140mM. Gray shaded regions are the two loops (89-93, 416-420). Black horizontal line is the standard deviation over contacts considering all four excipients. N-lobe and C-lobe iron coordinating residues are marked with black and red dashed lines, respectively.

**Figure S7.**
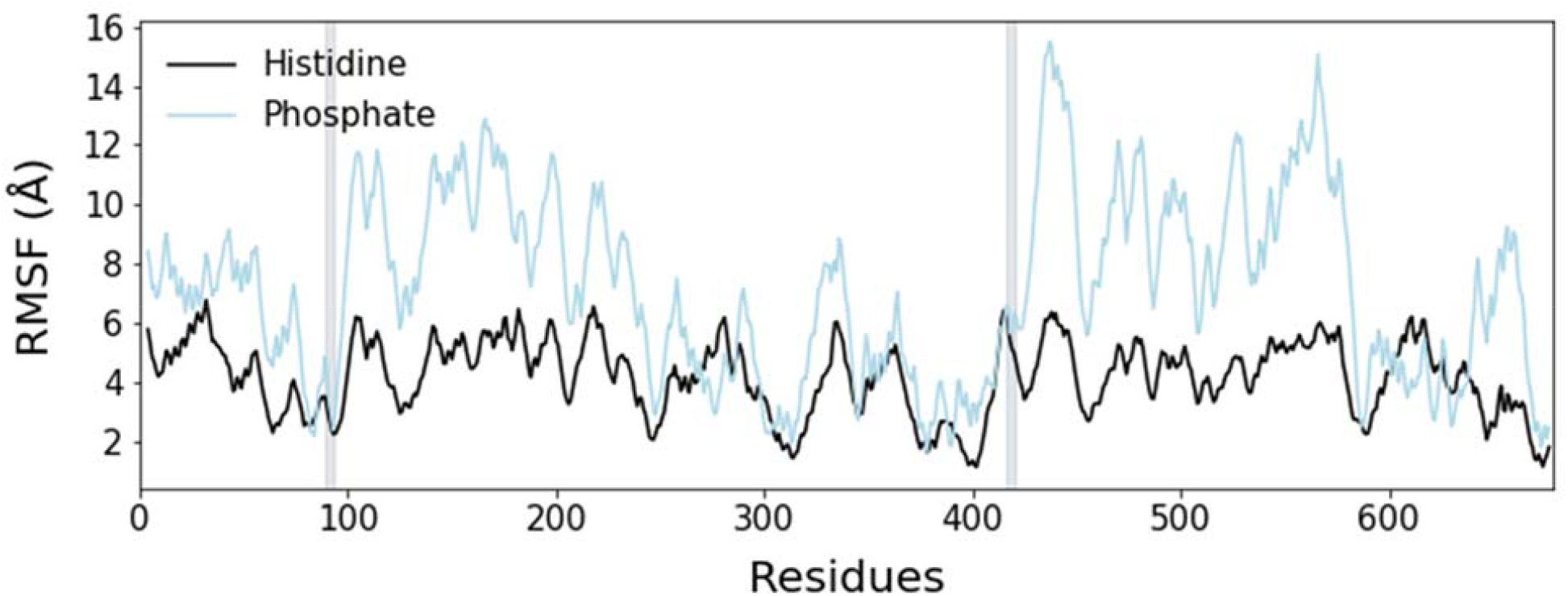
RMSF plot for the different excipient system at pH 6.5 for HO conformer. Gray shaded regions are the two loops (89-93, 416-420).

